# Signatures of antibiotic tolerance and persistence in response to divergent stresses

**DOI:** 10.1101/2023.02.05.527212

**Authors:** Huijing Wang, GW McElfresh, Nishantha Wijesuriya, Adam Podgorny, Andrew D. Hecht, J. Christian J. Ray

**Affiliations:** Center for Computational Biology, University of Kansas, 2030 Becker Dr., Lawrence, KS 66047 USA; Department of Molecular Biosciences, University of Kansas, 2030 Becker Dr., Lawrence, KS 66047 USA

## Abstract

In an environment with overly abundant lactose, a strain of the Gram-negative bacterium *E. coli* induces a persister-enriched phenotype and a heterogeneous pattern of growth rates. In high lactose conditions, the majority of cells are fast-growing, while a minority stochastically switch to a slow-growing, persister-prone phenotype that has higher ampicillin tolerance. Previously, bulk bacterial RNA-seq demonstrated broad changes in gene expression profiles for cells cultured in different lactose conditions, revealing multiple pathway regulatory regime switches enhancing its survivability for counteracting osmotic pressure in high lactose conditions with overflow metabolism.

We hypothesized that a set of unique gene regulatory signatures underlies antibiotic tolerance in the high lactose condition. To further understand the gene regulatory regime in slow-growing cells, the subpopulation of persister-prone cells was enriched with ampicillin treatment. The resulting culture was collected for transcriptomic analysis. The transcriptomic data were then analyzed for differentially expressed genes, signature genes, GO term enrichment, pathway enrichment, and flux balance analysis.

Our results show that under opposing stresses, the cells have similar and divergent responses. Cells exhibit upregulated assimilation pathways and downregulated biosynthesis pathways when encountering stresses. Post ampicillin treatment, cells in both high and low lactose conditions exhibit downregulated central metabolism to reduce growth. In the high-lactose concentration medium after ampicillin treatment, persisters may arise due to ferric imbalance-induced cell growth arrest and gene regulation due to *ssrA*-mediated downregulation-induced error-prone transcription.

## Introduction

Bacterial persisters were first discovered in 1944 by Joseph Bigger (1) by demonstrating that penicillin frequently failed to sterilize flask cultures of *Staphylococcus aureus*. Persisters usually enter dormancy or form biofilms to evade catastrophic environments and can resume growth after the lethal stress (2). Thus, persisters may cause recurrent chronic infections, such as lung fibrosis, and invalidate current antibiotics. Persisters are posing a serious challenge to contemporary medical treatment for bacterial infections. The bacterial killing curve typically presents as a biphasic killing pattern (3). Research into persister formation mechanisms has resulted in forming a tenuous consensus on several biologically important mechanisms including toxin-antitoxin (TA) systems and two-component systems in *E. coli* (4). Knocking out global regulators, such as DksA, DnaKJ, HupAB, and IhfAB, causes about a 10-fold decrease in persister formation (5). YgfA and YigB are candidate persister-related genes that change the nucleotide and metabolic dynamics inside cells (6). Many TA systems are identified as contributing to persister formation (7). Some TA modules contain the SOS-responsive Lex box regulatory motif, such as *symER*, *hokE*, *yafN*/*yafO* (7-14). We previously presented a novel, and counterintuitive, finding where both starvation and excess of lactose leads to a higher rate of persister formation in the *E. coli* B strain REL606 (15-17). This finding generalized the previous model where starvation stress drives environment-responsive persistence, but the mechanistic basis for these divergent stresses driving persistence have remained uncertain.

In the *E. coli* B strain model system, a robust cell wall confers survival in toxic quantities of lactose. Cells growing in high lactose conditions can stochastically switch to growth arrest, forming a persister-prone phenotype (17). As cell growth states are inheritable through lineage space, they form irregular clusters of growth-arrested cells under the microscope (16). In Part I of this study, we used bulk RNA-seq to identify putative mechanisms for persister formation, testing conditions representative of starvation, moderate lactose levels, and toxic conditions. We found that the toxified cultures exhibited multiple pathway regulatory mechanisms that enhanced their survivability under osmotic pressure and overflow metabolism (15). Bulk RNA-seq reveals population-level transcriptomic differences, where phenotypic heterogeneity is blended to give an average read in each condition. For instance, analysis in toxified conditions primarily captures the signal from fast-growing cells. To further separate the subpopulation of slow-growing cells, we performed a parallel set of experiments using ampicillin treatment with bulk RNA-seq performed on the surviving fraction of nominal persister cells. We hypothesized that a unique gene regulatory signature would underly the antibiotic tolerance in the high lactose condition.

Our results show that under opposite stresses, the cells have aspects of both similar and divergent responses. Cells exhibit upregulated assimilation pathways and downregulated biosynthesis pathways when encountering stresses. After ampicillin treatment, cells in both high and low lactose conditions exhibit downregulated central metabolism to reduce growth. Typically, in starvation, the persister arises due to stress response-induced growth arrest during antibiotic attack. In a medium with high lactose concentration, persisters may arise due to ferric imbalance induced cell arrest and epigenetic regulation due to *ssrA* downregulation induced error prone transcriptome environment.

## Results

### Differential gene expression analysis

To understand cellular responses to opposite stresses posed by differential lactose concentration, and the impact of this regulatory response on the subsequent ampicillin treatment response, we performed bulk bacterial RNA-seq for five conditions, including lactose 0.1 mg/ml (starvation) and lactose 50 mg/ml (toxification) with/without ampicillin treatment, and lactose 2.5 mg/ml (moderate) condition without ampicillin treatment. (Insufficient biomass survived antibiotic treatment for RNA-seq analysis in the moderate lactose concentration.) Significantly differentially expressed genes are defined with log_2_ fold changes > ±1 and FDR-adjusted p-value < 0.05. Six pairs of comparisons were performed for analyzing differentially expressed genes, as shown in Table 1. Data validation with PCA and clustering results are shown in Supplementary Figure S1.

**Table 1.** An overview of differential expression analysis results. Comparisons 1 to 6 are results from our RNA-seq data. The comparison column follows naming convention of “(comparison condition) – (reference condition)”. Thus comparison 1 to 4 have the same reference condition of moderate lactose, while comparison 5 and 6 are showing post-ampicillin treatment responses for different lactose conditions. For comparing condition, the numbers in parenthesis are lactose concentrations in the medium, and amp refers to ampicillin treatment.

| Comparison | Test condition<br>(Unit: mg/ml) | Reference condition<br>(Unit: mg/ml) | Total gene # | Upregulated gene # | Upregulated gene fraction | Downregulated gene number | Downregulated gene fraction |
| --- | --- | --- | --- | --- | --- | --- | --- |
| 1. Starving - moderate | lac(0.1) | lac(2.5) | 4490 | 1845 | 0.41 | 1145 | 0.26 |
| 2. Toxic - moderate | lac(50) | lac(2.5) | 4490 | 1755 | 0.39 | 830 | 0.18 |
| 3. Starving amp - moderate | lac(0.1) amp | lac(2.5) | 4490 | 2053 | 0.46 | 1053 | 0.23 |
| 4. Toxic amp - moderate | lac(50) amp | lac(2.5) | 4490 | 2129 | 0.47 | 1173 | 0.26 |
| 5. Starving amp - starving | lac(0.1) amp | lac(0.1) | 4490 | 1130 | 0.30 | 590 | 0.13 |
| 6. Toxic amp - toxic | lac(50) amp | lac(50) | 4490 | 1773 | 0.39 | 777 | 0.17 |

Differentially expressed gene (DEG) analysis generates a gene-level log_2_ fold change profile for each pair of comparisons in Table 1. Hierarchically clustering the log_2_ fold change profiles with Ward’s linkage separates different comparisons (Figure 1a). Comparing pre- and post-ampicillin treated condition, the toxic and starvation responses to ampicillin are most similar (the comparison 5 and 6 in Table 1), showing significant common responses facing ampicillin pressure. Comparing their transcriptional profiles to the moderate lactose condition, the toxicity DEG profile is notably most like starvation after ampicillin treatment, indicating the toxic phenotype state might share some persisters formation mechanism with starvation cells (Figure 1a). To understand how starvation-induced and toxicity-induced persister cultures differ, scatter plots for different DEG comparisons are shown in Figure 1b. We observe a strong positive correlation between the transcriptomic responses to ampicillin treatment for starving cells and toxified cells with the moderate lactose condition as the reference. This indicates the presence of common transcriptomic responses towards ampicillin stress, where the Pearson correlation is 0.89. A hyper-clustered region appears when comparing the pre- and post- ampicillin treatment conditions (Figure 1c): a high number of genes upregulated in ampicillin-tolerant cells, implying that either the global transcriptional profile adapts to ampicillin pressure, or that the persister cells are pre-adapted compared to the antibiotic-sensitive majority. The Pearson correlation for this comparison is 0.38. Still, there are significant DEGs that are uniquely differentially expressed for starvation persister and toxicity persister in both Figure 1b and 1c. We seek to understand the common and divergent persister formation mechanisms in opposite stress conditions.

**Figure 1.**
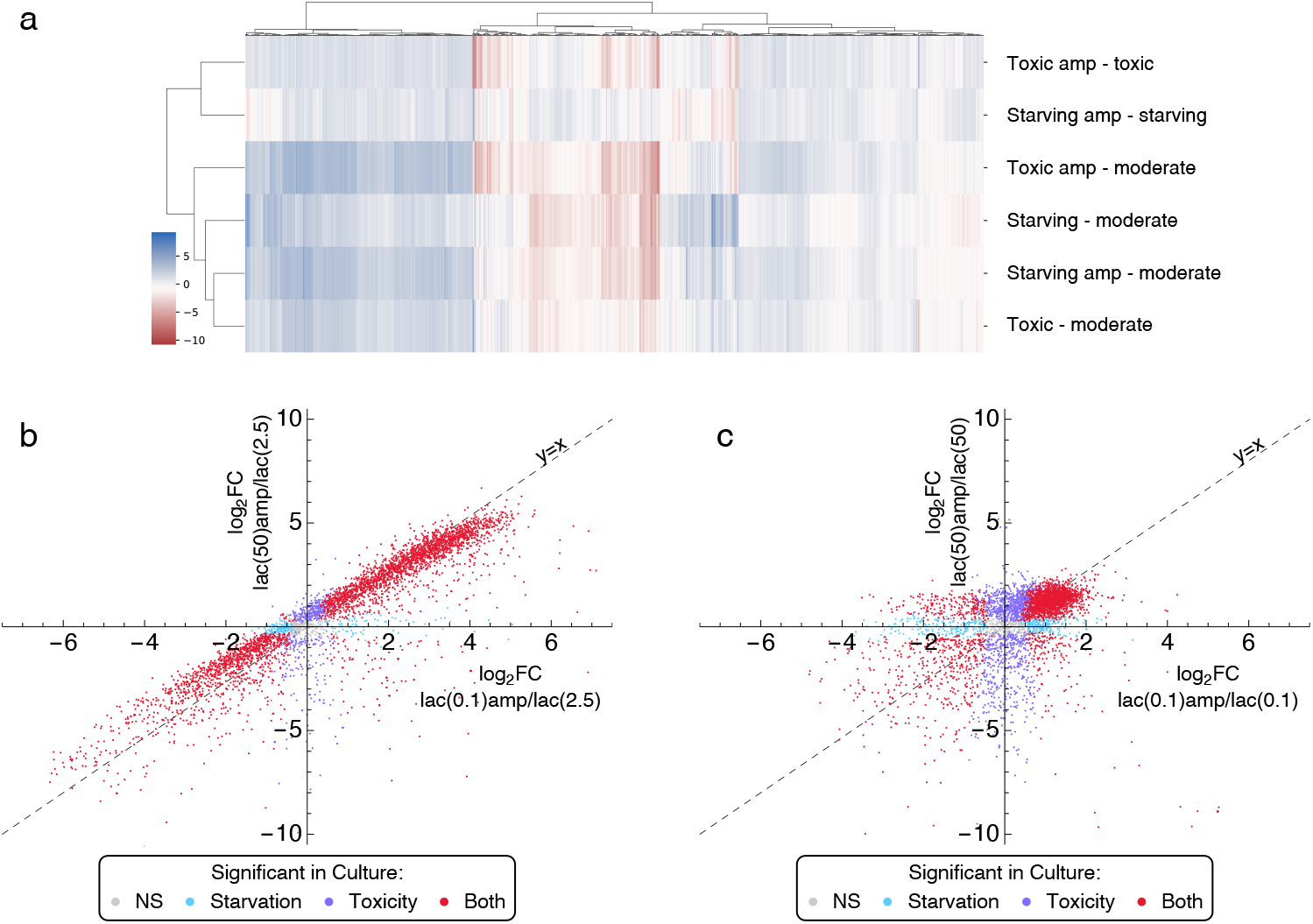
Differential expression analysis results for multiple comparisons. a. Differential expression log_2_ fold change clustering shows a closer similarity between the toxified cell transcriptomes and ampicillin-treated staving cell transcriptomes. b. Setting moderate lactose concentration as a reference, the transcriptomic responses to ampicillin treatment for starving cells and toxified cells are positively correlated with a Pearson correlation of 0.89. c. Comparing transcriptomic responses between post ampicillin treatment and non-ampicillin treatment, we find a large cluster in the first quadrant of the scatter plot, indicating the common responses for ampicillin stress in cells. The Pearson’s correlation between the axes is 0.38.

### *Lac* operon activities and growth regulator genes decreased after ampicillin treatment

Figure 2 explores persister-related gene expression responses after ampicillin treatment. In both starving and toxic conditions, we observe significantly downregulated lactose operon activities, while the non-functional *lacI* is not differentially expressed after ampicillin treatment (Figure 2b). The *lacI* gene in this strain carries a point mutation and is not able to perform binding-inhibition with the *lac* operon promoter, a choice driven by the desire to eliminate *lac* operon regulatory mechanisms as confounders in the analysis. Thus, any differential expression observed is caused by post-transcriptional mechanisms. Using the moderate lactose condition as the reference, post-ampicillin response of lactose operon downregulation in the toxic condition is qualitatively similar to, but stronger than, that of the starving condition (Figure 2a), with exception of *lacY*, the lactose permease. It has a more significant apparent reduction in expression in starvation conditions. This may be explained by selection for cells that randomly had higher permease permitting more robust survival in sub-optimal lactose concentrations.

**Figure 2.**
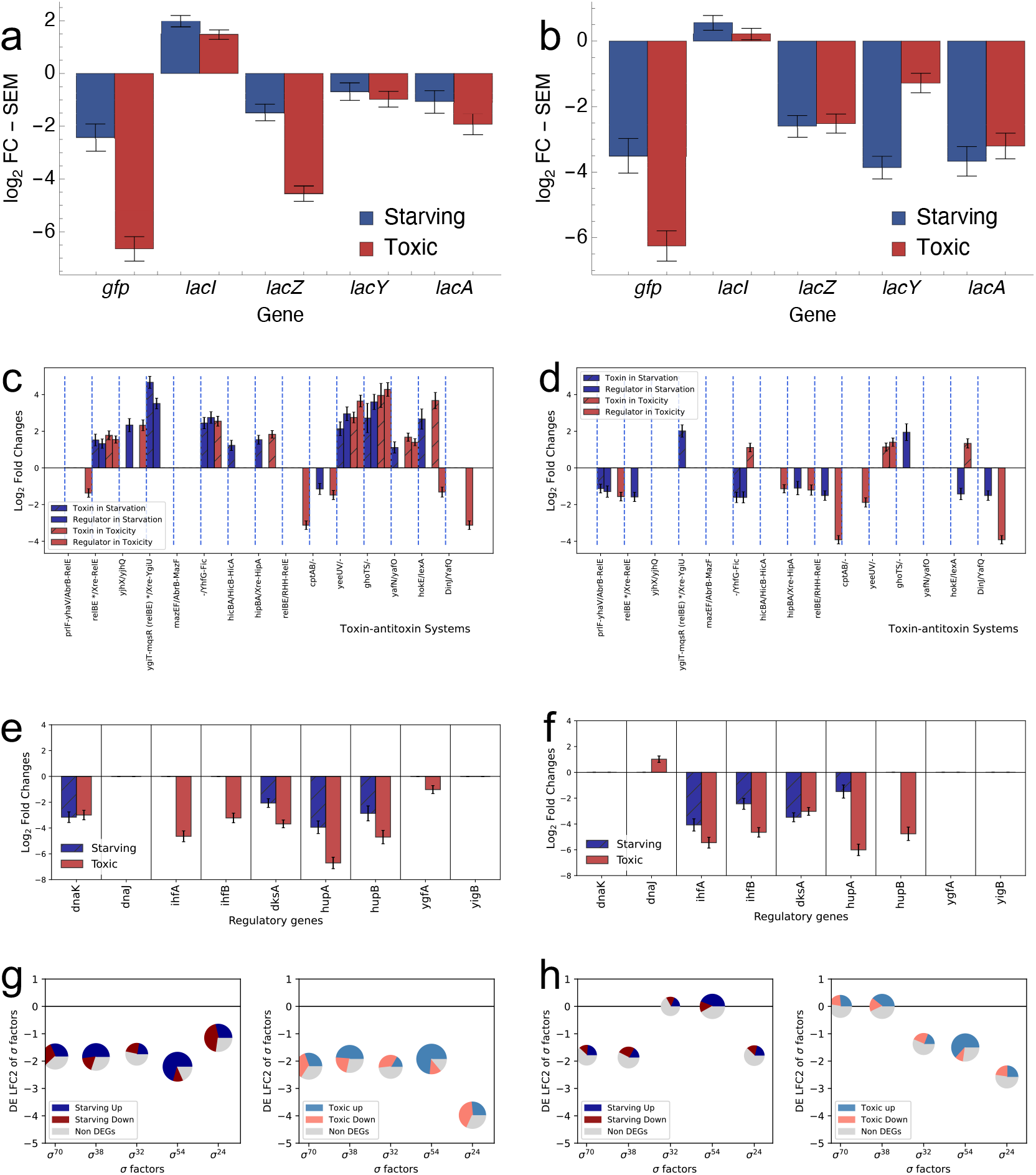
Differential expression log_2_ folder changes (LFC2) for genes of biological significance. a and b plots show lactose operon activities. c and d plots are showing LFC2 for all toxin-antitoxin pairs existing in REL606. e and f plots show LFC2 for biologically important global regulators. g and h are pie charts showing the LFC2 for sigma factors in REL606 and the fraction of genes that have been regulated by these sigma factors. a, c, e, and g are based on differential expression data analyzed with a reference level of moderate lactose condition, while b, d, f, and h compare post-ampicillin treatment responses to non-ampicillin treatment responses. Only significantly differentially expressed genes are plotted in Figure 2c-2f. Error bars, standard error of the mean (SEM).

Figure 2c, d show comparisons of 15 toxin-antitoxin systems. Compared to the moderate conditions, ampicillin treatment leaves cells with overall upregulation of toxin-antitoxin pairs in both starving and toxic cultures, with some exceptions, suggesting a major consistency for two components system induced persister formation (18, 19). Three toxin-antitoxin (TA) systems are upregulated in pairs under both toxic and starving conditions (Figure 2c), including *yeeUV* and *ghoTS*, which usually contribute to cell stasis and biofilm formation (20). *yeeV* is a toxin that inhibits cell division by targeting cytoskeletal proteins, including FtsZ and MreB (21). *ghoTS* is a novel type of TA module where the toxin GhoT is a membrane lytic peptide that causes ghost cell formation (lysed cells with damaged membranes), and GhoS is an antitoxin that cleaves the mRNA of the toxin. GhoT is known to induce persistency (22). Some TA modules have non-paired enrichment. *hipA* and *hokE* toxin expression and *cptB* antitoxin downregulation after ampicillin treatment in low and high lactose may further mediate the cell dormancy, host killing, growth arrest, or cell shape deformation (12, 23, 24). Despite these similarities between the low lactose persister and high lactose persister phenotypes, some TA modules perform differently. *mqsA* and *hicA* are enriched only in starvation, where *mqsA* triggers programmed cell death (25) and *hicA* is a toxin that cleaves mRNA and induces growth arrest (26). *yhaV* and *yafQ* are enriched only in high lactose persisters, where YhaV and YafQ are both ribosome-dependent mRNA interferase toxins (27, 28).

Global regulatory genes contributing to persister formation include 9 global growth regulators, for which LFC2 are shown in Figure 2e-2f (4). In ampicillin treatment, comparing to moderate lactose conditions, none of the growth regulators are upregulated. *dnaK*, *dksA*, *hupA*, and *hupB* are all downregulated in both types of lactose stress. Chaperone protein *dnaK* functions as a central hub in the *E. coli* chaperone network (29) with active participation in the response to hyperosmotic shock (30). *dskA* potentiates ppGpp regulation on stringently induced and repressed promoters (31). *hupA* and *hupB* are HU protein homologues that responsible for maintaining chromosome structure even in extreme environment (32, 33). *ihfAB* and *ygfA* are uniquely downregulated in ampicillin-treated high lactose conditions, suggesting lower DNA recombination activities and lower folate metabolic activities.

The sigma factor differential expression LFC2 is plotted in Figure 2g and 2h. Comparing to moderate conditions, ampicillin treatment seems to have a significant impact on the sigma factors, as all sigma factors in post ampicillin treatment conditions are downregulated, indicating globally downregulated transcription activity.

To summarize, we observed significant loss of expression of major regulatory genes responding to ampicillin treatment, such as growth regulator genes, and sigma factors, and general upregulation of toxin-antitoxin systems.

### Signature genes analysis reveal a unique set of genes that contribute to cell benefit in different condition

The bacterial gene regulatory mechanisms are not only a model system for understanding the fundamental regulatory principles for life but also crucial for developing a medical treatment for tackling health issues such as diarrhea, urinary tract infections, respiratory illness, pneumonia, and other illnesses. To infer cell state abundances from the transcriptional signature, we used CIBERSORTx, a computational framework for inferring cell type abundance and cell-type-specific gene expression (34). Our analysis yielded a subset of genes, where for each gene only one condition has significant expression, as shown in Figure 3a. Comparing Figure 1a and 3b, we found that clustering for all differentially expressed genes yields a similar dendrogram to clustering over the differentially expressed signature marker genes. Paired comparisons were performed and shown as heatmaps in Figure 3c-d, Figure S2–S3, and Table S1.

**Figure 3.**
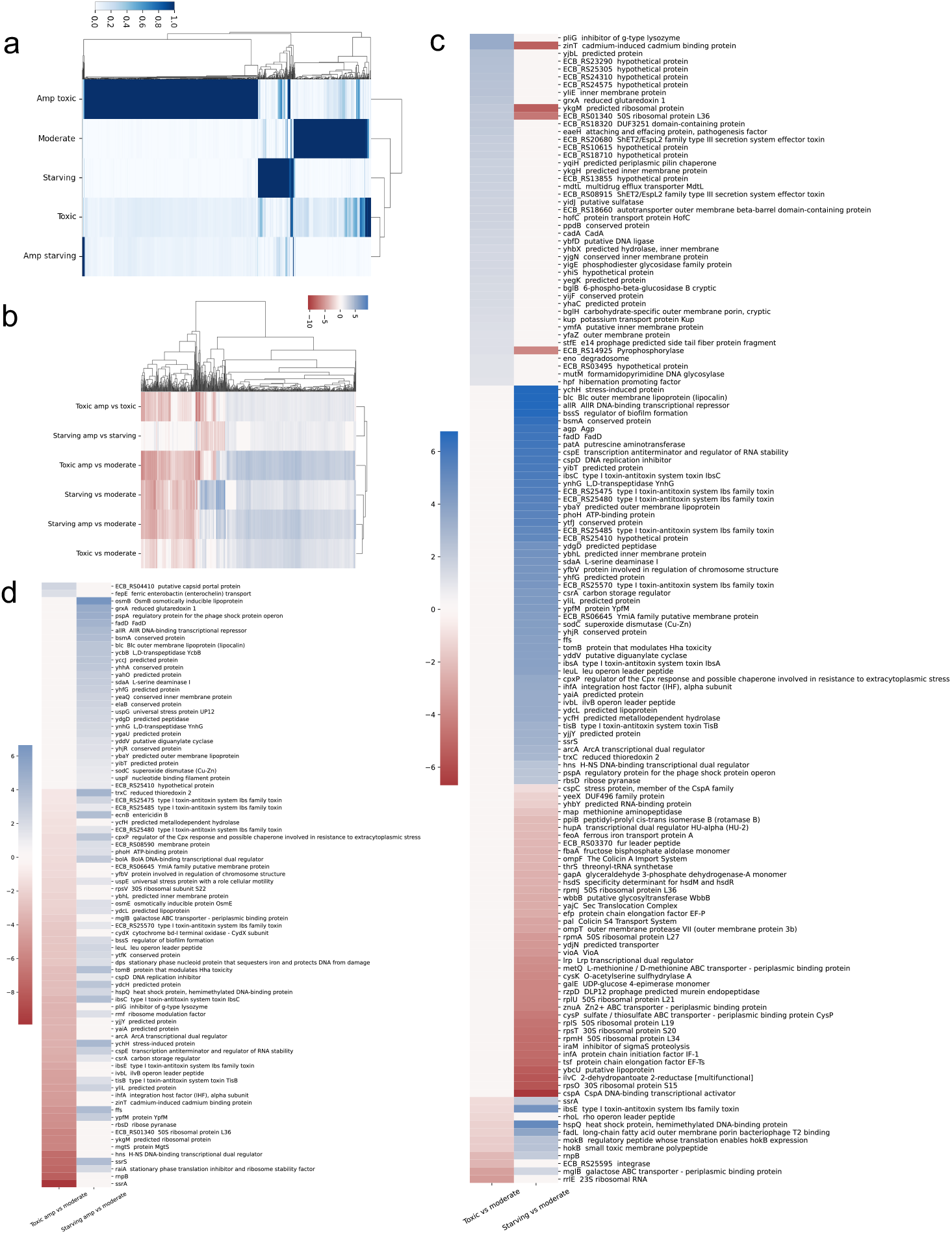
Differentially expressed signature marker genes were identified in different conditions. a. Signature marker genes are obtained with CIBERSORTx, and the clustered heatmap for signature marker gene expression for each condition is shown. The heatmap data is normalized by column. b. Differentially expressed signature marker genes are clustered in the heatmap. c and d show the clustered heatmap for differentially expressed and regulated signature genes in condition comparisons. If a conventional name for the sequence ID is found (Method 4.3), the conventional name is applied, otherwise, the sequence ID is shown.

We found significantly downregulated ribosomal proteins and ATP synthase genes, upregulated transporter, symporter, and membrane genes that are consistently regulated in opposite lactose stress conditions (Figure S2–S3). For lactose stress with no ampicillin treatment compared to the moderate condition, among 22 30S ribosomal subunit proteins, 19 are downregulated. S22 is the only upregulated 30S ribosomal subunit protein, which is typically abundant in stationary phase and can be activated by (p)ppGpp (35, 36). 27 out of 36 50S ribosomal proteins are downregulated in this condition. There are 57 inner or outer membrane proteins that are upregulated in stress conditions, while 6 are downregulated, indicating a consistent response in different stresses. 7 ATP synthase genes are downregulated, while 16 ATPase or ATP binding subunit genes are upregulated, explaining a tightening in energy production and heavier dissipation. 59 transporters are upregulated, while only *fecB*, the ferric dicitrate ABC transporter is downregulated. 7 of 8 symporters for Na^+^ and H^+^ are upregulated.

Compared to the moderate condition, we then examined differentially expressed genes that are regulated in different directions in opposite lactose stress conditions (Figure 3c). Interestingly, the signature genes identified many hypothetical proteins that do not have clear functionality, and that were enriched in toxic conditions. We also found membrane proteins enriched in toxic conditions, such as *yliE*, *yhbX*, and multidrug efflux transporter *mdtL*. Gene *mglB*, the galactose ABC transporter, is upregulated in starving condition and downregulated in toxic condition, suggesting that carbohydrate transport and metabolism is partially upregulated in starving conditions and downregulated in toxic conditions. As expected, in the starving condition, several toxin-antitoxin-related genes were upregulated, such as *ibsC*, *ibsA*, and *tisB*. We found that stress-induced protein *ychH*, ATP binding protein *phoH*, and carbon storage regulator *csrA* were upregulated, while there was downregulation of ribosomal proteins and DNA-binding transcriptional activator *cspA*. These pieces of evidence are consistent with a model where, in starvation, the cells tend to shut down transcription while activating toxin-antitoxin systems to inhibit growth.

Compared to moderate conditions, differentially expressed genes that are regulated differently in opposite lactose stresses after ampicillin treatment are shown in Figure 3d. We find *fepE*, the ferric enterobactin transport gene, is uniquely upregulated in the toxic ampicillin-treated condition, which may contribute to antibiotic tolerance (37). Different from starving ampicillin-treated conditions, some toxin-antitoxin systems are downregulated, such as *ibsC*, *ibsE*, and *tisB*. *ssrA* is significantly downregulated, which may effectively allow mutated transcripts due to low-fidelity transcription (38).

To summarize, by combining signature gene analysis with differential gene expression analysis, we were able to identify unique gene regulation in different conditions, compared to moderate, unstressed cultures. The analysis identifies many hypothetical proteins that may of biological importance for surviving the toxic condition and form persisters after antibiotic treatment.

### Enrichment analysis indicates upregulated transport activities in toxic conditions

We use CIBERSORTx (34) to isolate a subset of genes that determine cell phenotypes, i.e. the signature marker genes for each condition, and further imputed the cell fraction in each bulk RNA sequencing profile. Pseudo population fraction (Figure 4) shows a gradual shift of subtype transcript profiles as lactose concentration increases. The stressed cell population is less heterogeneous than that of non-stress conditions, while the stressed population shares some fraction of different stressed phenotypes across different stress conditions. For example, under starving conditions, about 15.9% of cells behave similarly to lactose starving cells under ampicillin treatment as presented in Figure 4a(i). Similarly, starving cells post ampicillin treatment (Figure 2d(i)) also contain 8.95% starving cells, 6.35% toxified cells, and 4.67% ampicillin treated toxified cells, with a minimum fraction of the non-stressed cell population. The cell population fraction analysis shows that the non-stress condition has the more heterogenous sub-phenotype, which makes sense since stress is usually a directional evolution pressure.

**Figure 4.**
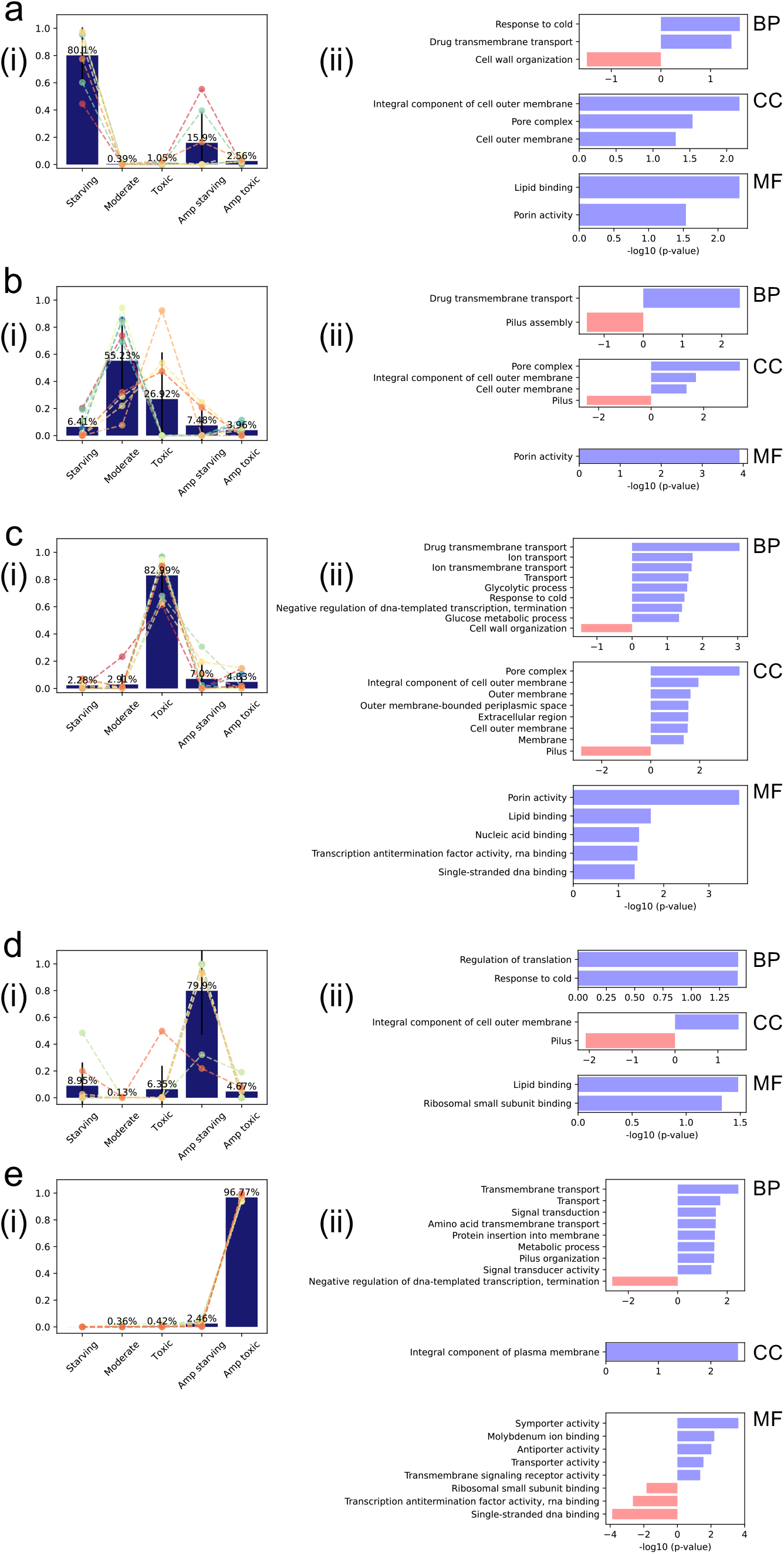
Panel (i) in a, b, c, d, and e is average imputed cell fractions based on the bulk mRNA-seq data for starving, moderate, toxic, starving post ampicillin treatment, and toxic post ampicillin treatment conditions. The scaler plot is imputed cell fraction for each transcription profile. We observe a gradually shifting cell fraction distribution for the transcription profiles in each condition. Panel (ii) is GO term enrichment for biological processes (BP), cell components (CC), and molecular function (MF).

Gene ontology (GO) is a formal representation for describing genes in classes, including molecular function, cellular component, and biological processes, and is widely used for understanding gene functionalities. The detailed process can be found in the method section.

For each condition, we applied GO term enrichment analysis for the signature gene matrix, as shown in Figure 4. The toxic and toxic post ampicillin treatment condition show significantly upregulated transport processes. Interestingly, in the toxic condition, there are upregulated central metabolism pathways, such as glucose metabolic process and glycolytic process, while also upregulating the transcription termination. In toxic post ampicillin treatment condition, the transcription is downregulated. In starving and starving post-ampicillin treatment conditions, we found the cold response is upregulated in both conditions.

We also performed pathway enrichment analysis on differentially expressed genes, shown in Figure S4. Consistent with our previous analysis (15), the pathway enrichment shows a major overlapping stimulons when responding to stress (Figure S4a). Common stress response pathways include upregulated assimilation processes, such as L-ascorbate degradation II, D-galactarate degradation I glycolate and glyoxylate degradation I. Common downregulated pathways include biosynthesis processes such as superpathway of purine nucleotides de novo biosynthesis II, superpathway of histidine, purine, and pyrimidine biosynthesis, superpathway of phenylalanine biosynthesis, superpathway of tryptophan biosynthesis. Interestingly, comparing the pre- and post-ampicillin treatment, we found downregulation for central metabolic pathways (Figure S4b).

### Flux balance analysis model predicts overflow metabolism in toxic conditions

Flux balance analysis is a framework for characterizing optimal flux distributions for a given objective reaction. We want to understand how different lactose concentrations impact the *in vivo* metabolic environment on our model system. The model is adapted from iECB-1328 by Monk et al (39), and the FBA model is constrained by both lactose influx and differentially expressed genes. Detailed procedures for FBA can be found in the Methods section. Briefly, we applied the constraints on FBA based on the naïve hypothesis that, for non-reversible reactions, upregulated genes promote reaction flux, thus constraining the lower boundary to a higher value, and vice versa (Figure 5a). We first optimized the moderate condition model, and then its fluxes were used for setting reaction flux boundaries in other conditions, based on fold changes of the differentially expressed genes.

**Figure 5.**
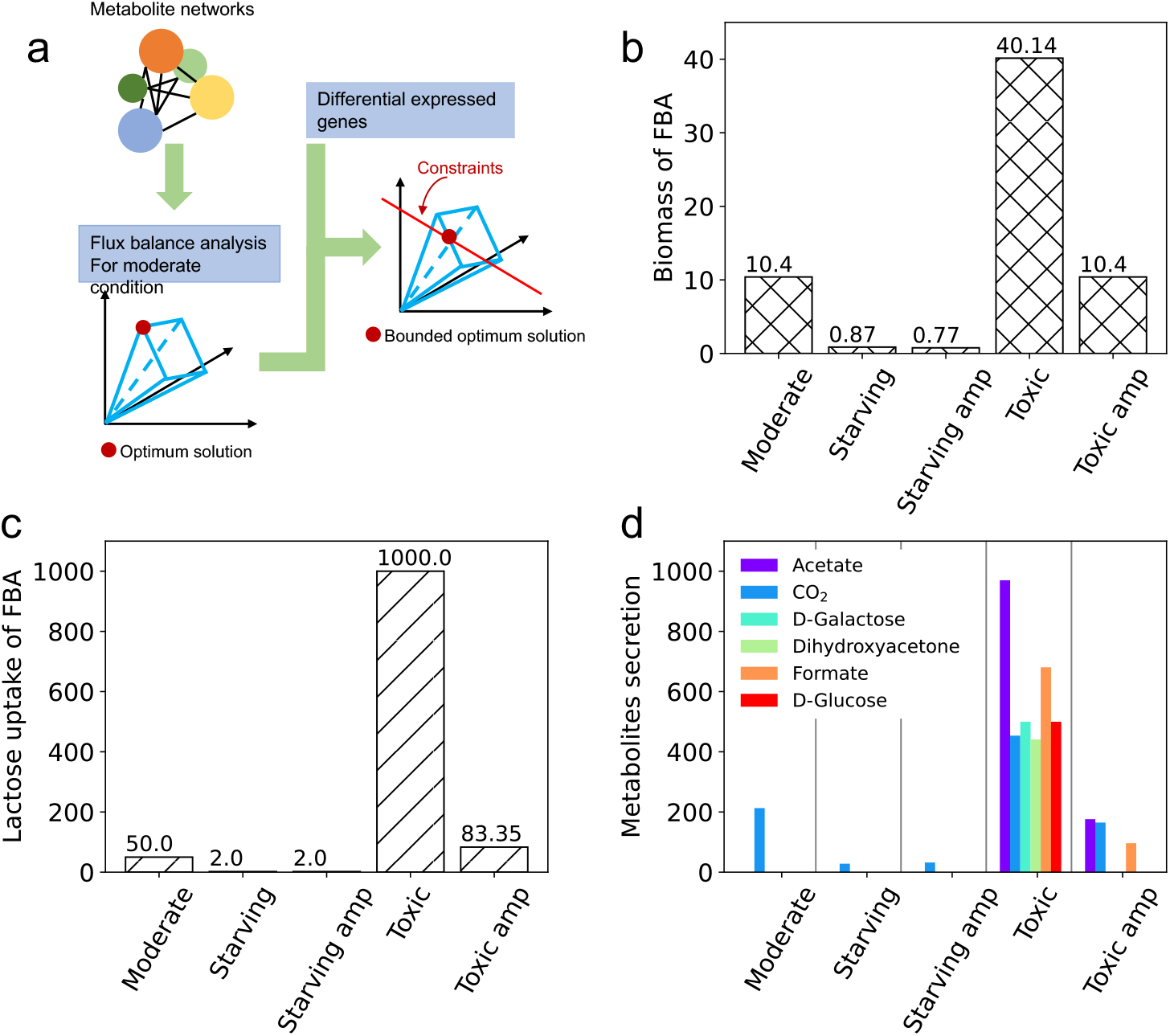
Flux balance analysis (FBA) shows metabolic benefit in different conditions. a. Differential expression analysis and FBA analysis is combined to locate an optimized solution for the model. We first optimize the FBA model for moderate condition and then use the optimized flux as boundary constraints based on the differentially expressed genes for other conditions. These constraints imposed by differentially expressed genes reduce the viable space for searching for optimized condition. b. Biomass was obtained from FBA in different conditions. The FBA model correctly predicted high and low growth rates in a toxic and starving condition. Interestingly, the FBA predicted that under ampicillin treatment, the cell cannot reach its peak biomass. c. Lactose consumption in different conditions. d. Predicted metabolite secretion under different conditions. In toxic condition, the model predicts overflow metabolism, such as secretion of acetate, and formate.

Optimizing the FBA model for different conditions, the solved optimized biomass is shown in Figure 5b. The FBA model correctly predicted low biomass in starving condition, and higher biomass in moderate and toxic condition. For moderate, starving, starving post ampicillin treatment, and toxic conditions, the limiting factor for biomass increase is the upper bound of the lactose uptake reaction. However, the limiting factor in toxic cultures after ampicillin treatment is gene regulation of lactose uptake. The predicted biomass in toxic ampicillin-treated cultures is dramatically decreased compared to the toxic condition, and its lactose uptake and metabolite secretion is less than toxic condition (Figure 5c-d). Thus, we can infer that cell growth is hindered by gene regulation, which reduced lactose uptake into the metabolic pathways.

FBA predicts overflow metabolism in toxic conditions both with and without ampicillin treatment. In toxic conditions, cells secrete the highest fluxes for acetate, formate, and CO_2_ compared to all other conditions. To understand these dynamics, we specifically explored central metabolic pathways, as shown in Figure S5–S6. Under the moderate condition (Figure S5e), the highest flux occurs in glycolysis, and the TCA cycle, pentose phosphate pathway, and anaplerotic pathway have fluxes near median of all fluxes in the central metabolism. However, this is the opposite in starving conditions either with or without ampicillin. This is probably due to the TCA cycle being the main energy-producing pathway. The absolute value for fluxes in starving conditions (with or without ampicillin) is small, thus limiting its biomass gain in those conditions. Comparing to the toxic ampicillin-treated condition, the cultures in toxic conditions without ampicillin have extremely high lactose uptake (Figure S5c). These results are consistent with pathway enrichment analysis results, where comparing the ampicillin-treated with no treatment conditions, central metabolism is decreased for both starving and toxic conditions (Figure S4b).

## Discussion

Transcriptomic data is a powerful tool for us to understand cell phenotypes. In this paper, we applied gene differential expression analysis, signature gene analysis, GO term enrichment analysis, pathway enrichment analysis, and flux balance analysis to compare cell responses to opposite lactose stress.

In previous research, we found that *E. coli* B REL606 cells grown in high lactose condition exhibit random growth arrest and can form more persisters after ampicillin treatment, compared to low lactose culture condition (16, 17, 40). We hypothesized that the persister formation in high lactose condition is due to epigenetic alterations, where in high lactose concentration environment induce dynamic changes to the transcriptome and subsequently forms new phenotypes. Typically, persisters formed due to *E. coli*’s stress response, such as activation of toxin-antitoxin systems and stress response systems. Here we show that persisters that arise in medium with high lactose concentration have different pathway mechanisms and metabolism for antibiotic tolerance.

There are similarities when cells respond to lactose stress. The pathway enrichment analysis shows that in all four stressed conditions, including toxic, toxic post-ampicillin treatment, starving, and starving post-ampicillin treatment conditions, many assimilation pathways were upregulated, and many biosynthesis pathways are downregulated. The upregulated assimilation pathways include L-ascorbate degradation II, arginine degradation II, D-galactarate degradation I, and so on (Figure S4a). The downregulated biosynthesis pathways include superpathway of purine nucleotides de novo biosynthesis II, superpathway of histidine, purine, and pyrimidine biosynthesis, superpathway of phenylalanine biosynthesis, superpathway of tryptophan biosynthesis, and so on. In both toxic and toxic post ampicillin treatment conditions, GO terms for transporter are enriched. In both starving and starving post ampicillin treatment conditions, a GO term for cold response is enriched, and toxin-antitoxin systems are activated. After ampicillin treatment, the opposite stress responses both include downregulation of *lac* operon, central metabolism, growth regulators and sigma factors.

The signature gene analysis, GO term enrichment, and flux balance analysis reveals distinct cellular response in different conditions. In toxic condition, we found genes for membrane structure, and efflux transporter are upregulated, and genes for galactose binding are downregulated. In GO term enrichment for the toxic condition, we found an interesting result that both central metabolism and transcription termination are enriched, probably reflecting the fact that growth arrest is the result of such regulation. Flux balance analysis predicts overflow metabolism in toxic conditions, consistent with previous research. In starving conditions, there is a significant decrease in ribosomal proteins and DNA transcription activator genes. In toxic post-ampicillin treatment, transcription molecular function terms are downregulated. The signature gene analysis reveals unique upregulation of ferric enterobactin transport *fepE* and significant downregulation of *ssrA*. Iron homeostasis is important for maintaining the proper concentration of reactive oxygen species (ROS) inside the cell (41). An overabundance of ferric iron is typically toxic to cells (42). The unique upregulation of *fepE* may introduce toxicity for the cell to enter growth arrest, thus forming persisters. *ssrA* is a remarkable RNA molecule that can mark incomplete proteins for degradation (43). Significant downregulation of *ssrA* might permit tolerance of proteins targeted to proteosomes that contribute to non-genetic regulation. Different from the persisters that arise in starvation, the persisters in toxic post-ampicillin treatment also show down regulation of several toxins. Interestingly, our results also show many hypothetical proteins as uniquely regulated in toxic and toxic post-ampicillin treatment conditions, which may be genes of interest in future research.

## Methods

Our model system uses the *E. coli* strain B REL606, which has a unique enriched survival profile in toxic lactose concentrations compared to the K-12 strain (44). This difference may arise from a more robust cell wall and has been shown to increase the survival of this strain compared to others in ionically and osmotically suboptimal media (44-46).

### 4.1. Persister Enriched RNA-Seq

An E. coli REL606 *lacI*^−^ strain transformed with Tn7∷PlacO1GFP(KanR) used in previous experiments in the lab (44, 47) was inoculated in LB medium from a −80°C bacterial stock and grown for 16 hours in a 37°C shaking incubator. The LB culture was then resuspended (1:1000) into 5mL of Davis Minimal medium (DM; Difco) supplemented with thiamine and one of three lactose concentrations (0.1 mg/mL, 2.5 mg/mL, and 50 mg/mL). The cultures were allowed to acclimatize for 24 hours before being resuspended (1:1000) into 5mL of the same Davis Minimal medium and lactose concentration. Cultures were grown long enough (8 hours in 2.5 mg/mL lactose, 10 hours in 50 mg/mL lactose, and 12 hours in 0.1 mg/mL lactose) to provide enough biomass for RNA-Seq after antibiotic treatment. After the initial growth phase, 1.5 mL of untreated (no antibiotic) cell culture was RNA isolated according to the RNA isolation procedure below, dosed to 50 μg/mL concentration of ampicillin, and incubated for 24 hours in a 37 °C shaking incubator for persister cell enrichment. 3.0 mL of persister-enriched culture was then RNA isolated (procedure below) for lactose concentrations 0.1 mg/mL and 50 mg/mL. Lactose concentration 2.5 mg/mL required large experimental deviations to achieve enough post-ampicillin biomass requirements and was omitted.

### 4.2. RNA Sequencing

Cell cultures were pelleted in a microcentrifuge (10,000 G for 2 minutes), washed in PBS buffer twice, and resuspended in 500 μL of RNA-Later (ThermoFisher) and stored in at −20°C for one week. The persister-enriched 50 mg/mL lactose culture was unable to be preserved by RNA-Later and proceeded to RNA isolation immediately. RNA isolation was performed using Direct-zol (Zymo) and TRIzol reagent (ThermoFisher) and stored in a −80°C freezer overnight. Isolated RNA was ribo-depleted by RiboZero (Illumina) using ethanol washing to precipitate the RNA. Library preparation was completed using NEBNext Ultra II Directional RNA Library Prep Kit for Illumina (New England Biolabs) and sequenced using MiSeq v3 Paired End 150 bp (Illumina).

### 4.3. Genome re-annotation with Ecocyc and RegulonDB

The RNA transcript quantification was performed by kallisto (44) using genome NC_012967.1 (48) (*Escherichia coli* B str. REL606) was used as the reference genome. Kallisto was run using paired-end data and 10 bootstrap samples. As there is a recent update to B REL606 sequence on NCBI, sequence offsets exist between Ecocyc annotations and NCBI sequence annotations for the B REL606 strain. Thus, annotations from different databases were recognized by pairing annotation with gene locus. Furthermore, lacking extensive annotation of gene regulation in REL606, we used the K-12 MG1655 strain annotation based on gene and gene product similarities. Thus the annotation crossing strains are mapped by sequence alignment (Supplementary Methods), where databases Ecocyc (49) and RegulonDB (50) were combined for further analyses.

### 4.4. Differential Expression Analysis

We used R package DESeq2 for gene differential expression analysis. The RNA transcription quantification data were firstly clustered for isolating outliers using principal component analysis (PCA) (Figure S1a). Two samples were taken out from subsequent analysis due to wrong clustering in the PCA plot with a high number of missing values. The rest samples were regarded as reliable and went into the DESeq2 pipeline. As a verification of sample treatment, a hierarchical clustering using Wald significance test is performed on all samples, and the sample treatment was retrieved perfectly (Figure S1b).

DESeq2 pipeline includes size factor estimation, dispersion estimation, and DEG tests. Low-count RNA quantifications are noisy and may decrease the sensitivity of DEGs detection (51). Size factors were calculated with a subset of control genes, which are non-regulated genes according to RegulonDB (50) and have expression higher than threshold 10 across all replicas. RNA sequencing profiles obtained from lactose starvation (0.1 mg/ml), moderate (2.5 mg/ml) and toxification (50 mg/ml) conditions were analyzed. Setting transcriptome quantification from the moderate lactose condition, we applied the adaptive-T prior shrinkage estimator “apeglm” and used Wald significance tests for detecting DEGs and obtaining the log_2_ fold changes (LFC2).

### 4.5. CIBERSORTx signature gene analysis and cell fraction deconvolution

CIBERSORTx is a machine learning frame for analyzing signature genes for phenotype transcriptome profiles and estimating cell type abundance from bulk transcriptomes. To generate the signature genes, the raw count profile was used for analyzing significant gene signatures for each phenotype. Then identified signature matrix file and the raw counts profile were used to impute cell phenotype fractions.

### 4.5. Enrichment analysis by FGSEA

The signature gene matrix is obtained with CIBERSORTx. The GO terms are obtained from the EcoCyc database. The GO term enrichment analysis is then performed with the R package FGSEA (52). The significant threshold is set to a p-value < 0.05.

There are total of 368 pathways in Ecocyc for B REL606. There are 6 pathways doesn’t involve gene products, thus we are discussing the 362 pathways that include enzymatic reactions. Pathway enrichment is analyzed by R package FGSEA (52). Differential expressed genes are pre-ranked by their log_2_ fold change. Pathway gene sets are defined by referring to the EcoCyc database. The pre-ranked gene data and pathway gene sets are then going through the fgseaMultilevel function for obtaining final results. The minimum gene set size is set to 3, and the bootstrap number is set to 200. The cutoff threshold for pathway significance adjusted p-value is 0.05.

### 4.6. Flux balance analysis

The metabolite benefit was assessed using flux balance analysis (FBA), with metabolite flux bounding conditions constrained by differentially expressed genes. Firstly, we prepared the model by updating gene IDs, for keeping consistent with our sequencing results. The model we used is iECB-1328 by Monk *et al* (39), including 1951 metabolites, 2748 reactions, and 1329 genes. Secondly, setting reaction BIOMASS_Ec_iJO1366_core_53p95M as the objective reaction, the FBA was run for the moderate lactose condition. The model fluxes are kept as a reference. Lastly, combining differential expression analysis results and FBA on various conditions were performed. We used the differential expression analysis results for adjusting the reaction bounding condition. To set reliable constraints, we set a higher DEG threshold for log_2_ folder change to 4. For a non-reversible reaction regulated by a DEG, if the DEG is upregulated, we reset the lower bound of the reaction to the flux obtained in moderate condition. If the DEG is downregulated, we reset the upper bound of the reaction the flux obtained in moderate condition, reflecting the regulatory effect on the metabolic flow.

### 4.7. Code availability

Analysis code is available at https://github.com/jcjray/ecoli_divergent_stress_pipeline

---

**Figure S1.**
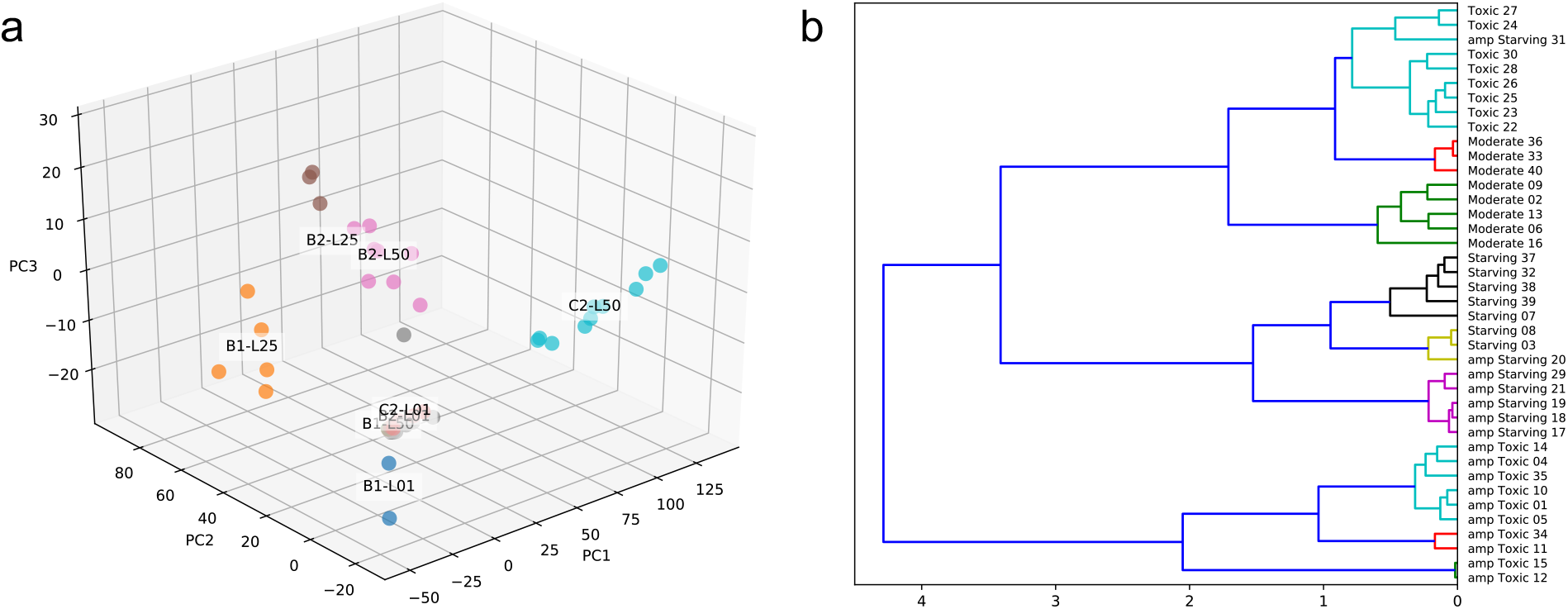
Data quality check results. a. PCA graph of first three dimensions for raw transcription profile. b. Normalized replica transcriptomic profile clustering restores the replica conditions.

**Figure S2.**
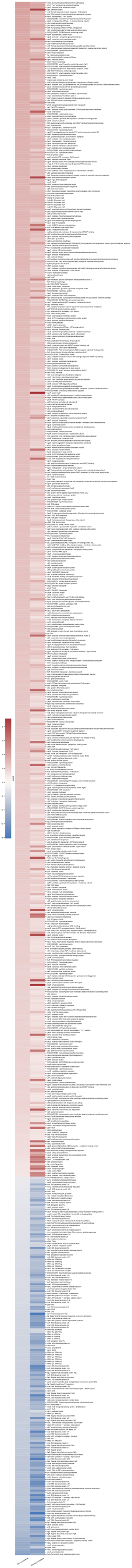
Signature genes that were regulated in the same direction in the opposite lactose stress compared to moderate condition.

**Figure S3.**
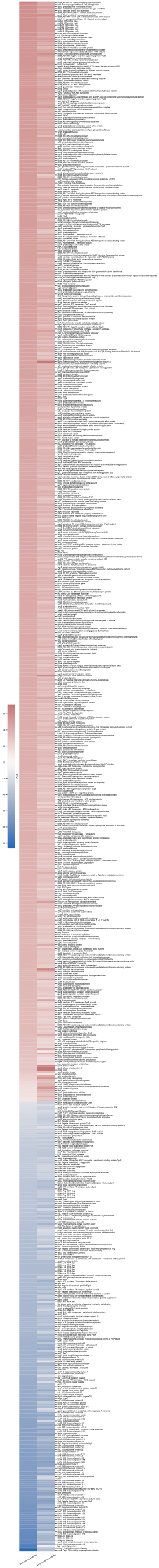
Signature genes that were regulated in the same direction in the opposite lactose stress post ampicillin treatment compared to moderate condition.

**Figure S4.**
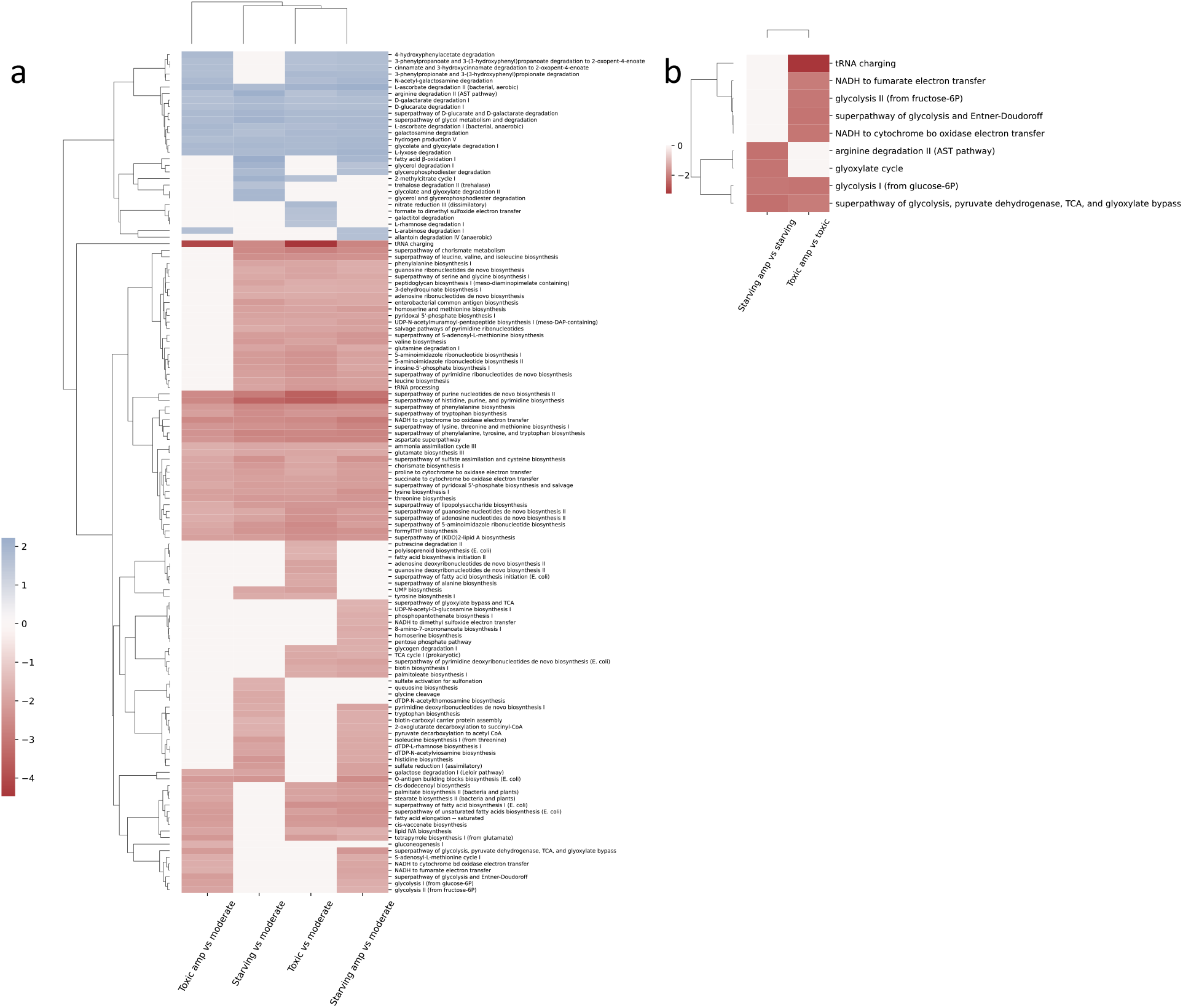
Pathways enrichment score clustering shows overlapping respond stimulons. a. Enriched pathways for toxic, toxic post ampicillin treatment, starving and starving post ampicillin treatment conditions compared to moderate condition. b. Enriched pathways for pre- and post-ampicillin treatment condition comparisons.

**Figure S5.**
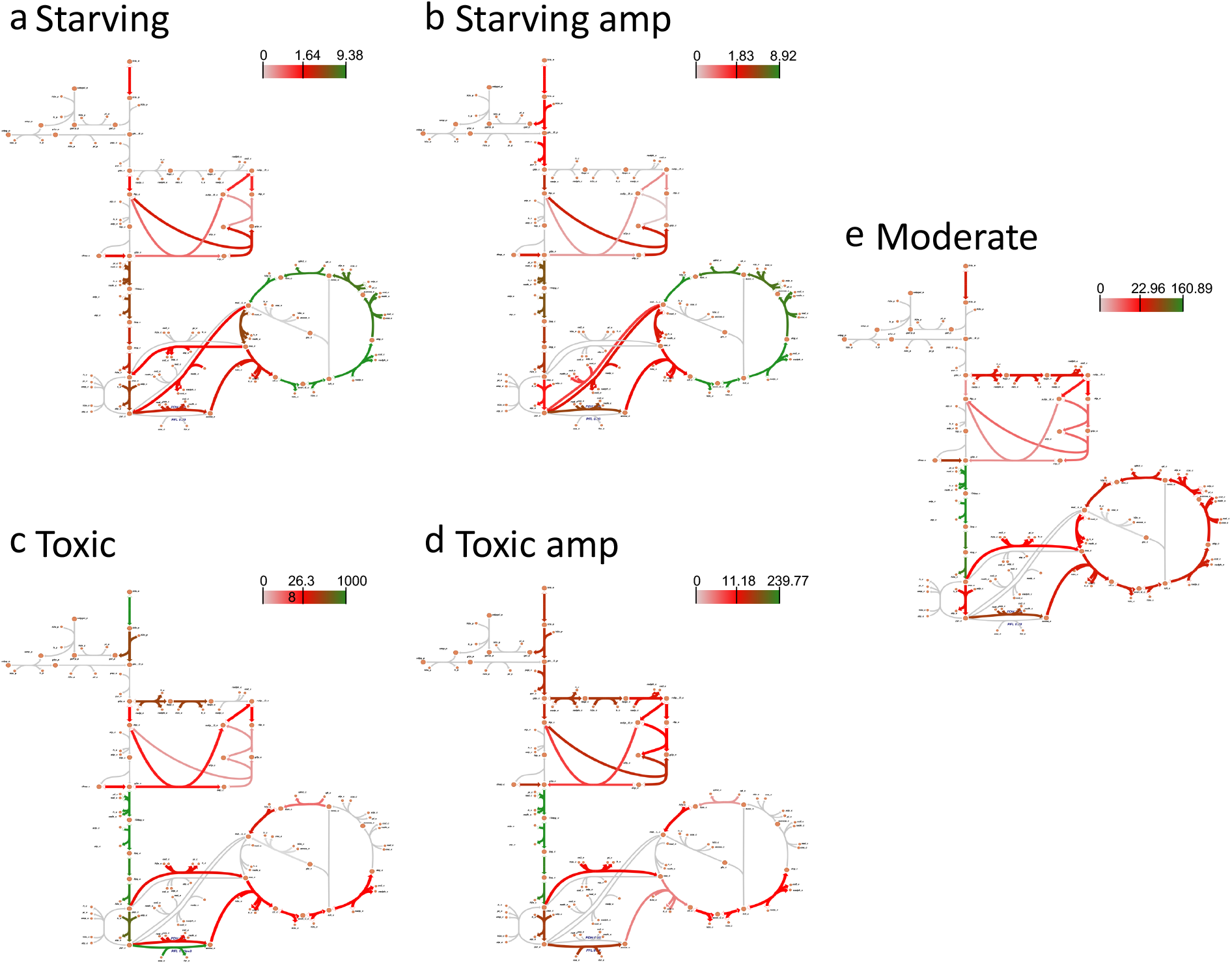
Fluxes in central metabolism are mapped for different conditions. The grey paths are low flux paths, the red paths are median fluxes, and the green paths are of the highest fluxes.

**Figure S6.**
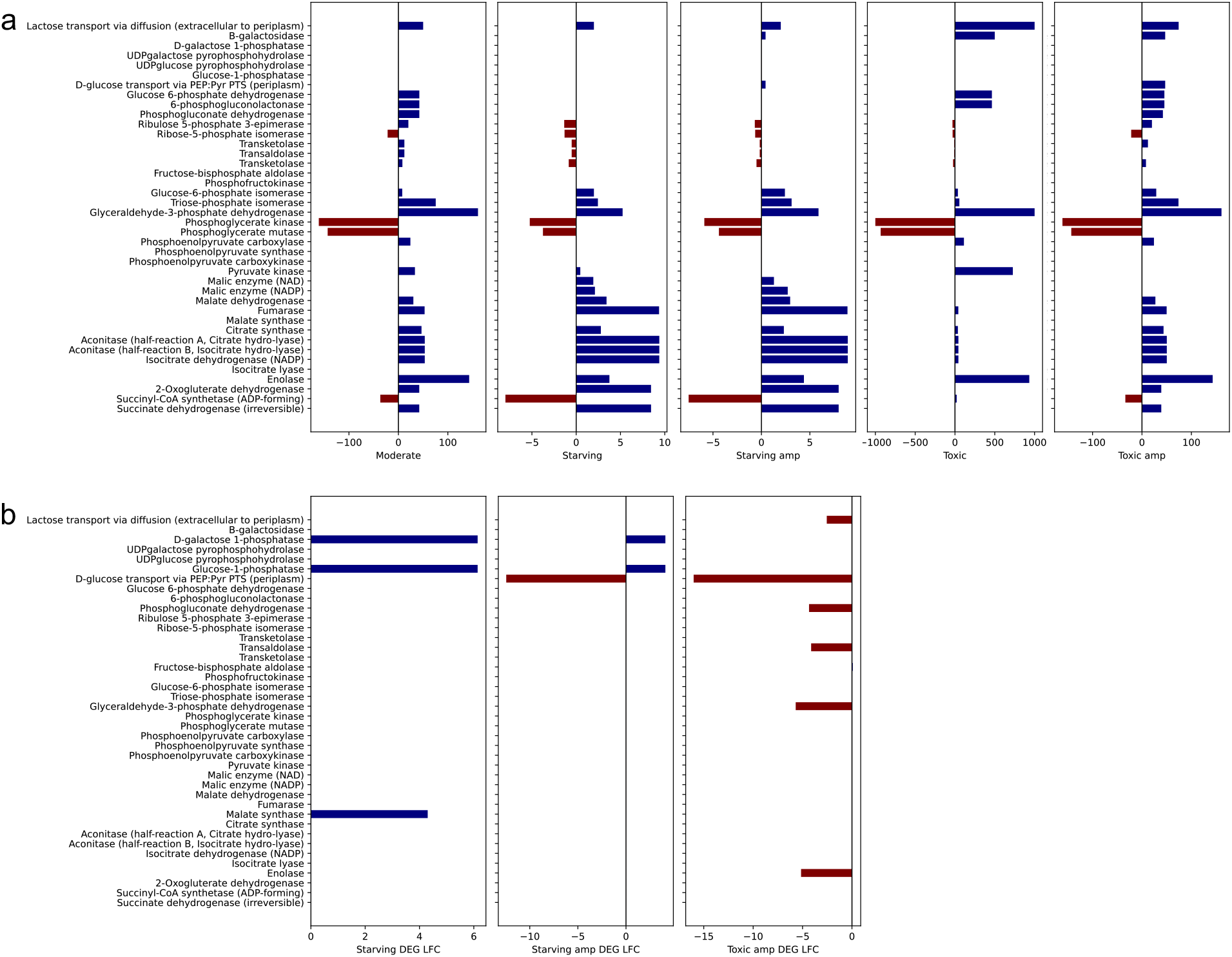
Flux balance analysis (FBA) results for central metabolism. a. Central metabolism reaction fluxes in each condition. Positive fluxes (blue) indicate that the reactions happen in the forward direction, while negative fluxes (red) indicate that the reactions happen in the reverse direction. b. The differentially expressed genes that were used for constraint of the FBA model.

**Table S1.** Overview of signature genes that are differentially expressed in paired comparisons.

| Differential expression (DE) analysis pairs | DE pair 1<br>(Unit: mg/ml) | DE pair 2<br>(Unit: mg/ml) | Total gene # | Same regulatory direction |  | Subtotal DE gene # | Different regulatory direction |  | Subtotal DE gene # |
| --- | --- | --- | --- | --- | --- | --- | --- | --- | --- |
|  |  |  |  | Upregulated gene # | Downregulated gene # |  | Upregulated signature gene #<br>(DE pair 1/2) | Downregulated signature gene #<br>(DE pair 1/2) |  |
| 1 Post amp treatment vs moderate | lac(50) amp vs lac(2.5) | lac(0.1) amp vs lac(2.5) | 4490 | 536 | 219 | 755 | 2/58 | 54/0 | 82 |
| 2 None amp treatment vs moderate | lac(50) vs lac(2.5) | lac(0.1) vs lac(2.5) | 4490 | 518 | 167 | 685 | 45/10 | 58/44 | 146 |
| 3 Post amp vs pre amp | lac(50) amp vs lac(50) | lac(50) amp vs lac(50) | 4490 | 417 | 78 | 495 | 101/172 | 36/43 | 316 |

